# Perforation of the host cell plasma membrane during *Toxoplasma gondii* invasion requires rhoptry exocytosis

**DOI:** 10.1101/2024.10.12.618018

**Authors:** Frances Male, Yuto Kegawa, Paul S. Blank, Irene Jiménez-Munguía, Saima M. Sidik, Dylan Valleau, Sebastian Lourido, Maryse Lebrun, Joshua Zimmerberg, Gary E. Ward

## Abstract

*Toxoplasma gondii* is an obligate intracellular parasite, and the delivery of effector proteins from the parasite into the host cell during invasion is critical for invasion itself and for parasite virulence. The effector proteins are released from specialized apical secretory organelles known as rhoptries. While much has been learned recently about the structure and composition of the rhoptry exocytic machinery and the function of individual rhoptry effector proteins that are exocytosed, virtually nothing is known about how the released proteins are translocated across the host cell plasma membrane. Previous electrophysiology experiments reported an unanticipated observation that invasion by *T. gondii* is preceded by a transient increase in host cell plasma membrane conductance. Here, we confirm this electrophysiological observation and propose that the conductance transient represents a parasite-induced perforation in the host cell plasma membrane through which rhoptry proteins are delivered. As a first step towards testing this hypothesis, and to provide higher throughput than patch clamp electrophysiology, we developed an alternative assay to detect the perforation. This assay utilizes high-speed, multi-wavelength fluorescence imaging to enable simultaneous visualization of host cell perforation and parasite invasion. Using this assay, we interrogated a panel of mutant parasites conditionally depleted of key invasion-related proteins. Parasites lacking signaling proteins involved in triggering rhoptry secretion (*e.g.*, CLAMP) or components of the rhoptry exocytic machinery (*e.g.*, Nd9, RASP2) are defective in their ability to induce the perforation. These data are consistent with a model in which the perforating agents that disrupt host cell membrane integrity during invasion – and may thereby provide the conduit for delivery of rhoptry effector proteins – are stored within the rhoptries themselves and released upon contact with the host cell.

## Introduction

Invasion of host cells is a critical step in the lytic cycle of the protozoan parasite *Toxoplasma gondii* and essential for establishing infection. The overall process of invasion is well conserved among apicomplexan parasites, including *T. gondii* and the related parasites that cause malaria (*Plasmodium* spp.) and cryptosporidiosis (*Cryptosporidium* spp.). Invasion involves the recognition of and attachment to a host cell, delivery of parasite proteins into that host cell, and parasite-driven internalization. Prior to invasion, the proteins to be translocated into the host cell are stored in the rhoptries, which are club-shaped exocytic organelles that consist of a posterior bulb and an elongated neck docked at the apical end of the parasite. Delivery of rhoptry proteins into the host cell is essential for invasion. For example, translocated rhoptry neck proteins (RONs) form a complex at the host cell plasma membrane that serves as a binding site for the parasite, providing traction at the “moving junction” through which the parasite propels itself into the host cell during invasion (Alexander, Mital, Ward, Bradley, & Boothroyd, 2005; Besteiro, Michelin, Poncet, Dubremetz, & Lebrun, 2009; Lebrun et al., 2005). Translocated rhoptry bulb proteins (ROPs) act downstream of invasion to suppress the innate immune response and manipulate the host cell in ways that facilitate expansion of the parasite population (Butterworth et al., 2022; Fukumoto et al., 2021; Hernández-de-Los-Ríos et al., 2019; Kochanowsky, Thomas, & Koshy, 2021; Li et al., 2020; Steinfeldt et al., 2010). Given their key biological functions, it is not surprising that rhoptry effector proteins are crucial for parasite virulence (Saeij et al., 2006; Shwab et al., 2016; Taylor et al., 2006).

The rhoptry exocytic machinery has become increasingly well defined in recent years. A search for apicomplexan orthologs of genes important for the discharge of exocytic organelles in ciliates, fellow members of the Alveolate superphylum, has identified a growing set of proteins important for rhoptry exocytosis (Aquilini et al., 2021; Sparvoli et al., 2022). Through reverse genetics and *in situ* cryo-electron microscopy, the detailed structures of the rhoptry secretory apparatus (RSA) and many of its molecular components have been determined in *Toxoplasma*, *Plasmodium*, and *Cryptosporidium* (Aquilini et al., 2021; Bisson, Hecksel, Gilchrist, Carbajal, & Fleck, 2021; Gui et al., 2022; Mageswaran et al., 2021; Martinez et al., 2022; Segev-Zarko et al., 2022; Sun et al., 2024). The RSA lies at the extreme apical tip of the parasite plasma membrane and forms the apical rosette, a transmembrane structure to which an underlying membrane-bound apical vesicle (AV) is typically docked (Aquilini et al., 2021; Paredes-Santos, de Souza, & Attias, 2011; Porchet-Hennere & Nicolas, 1983). The AV links the RSA to the tips of the rhoptries (Aquilini et al., 2021; Mageswaran et al., 2021). Exocytosis of lumenal rhoptry proteins through the rosette therefore involves two membrane fusion events: the rhoptry tip with the AV, and the AV with the parasite plasma membrane (Fig. 1A). The order in which these fusion events take place and whether they are independent or co-regulated is currently unknown, although recent ultrastructural data from *Plasmodium* merozoites suggest that rhoptry-to-AV fusion occurs first (Martinez et al., 2022).

**Figure 1.**
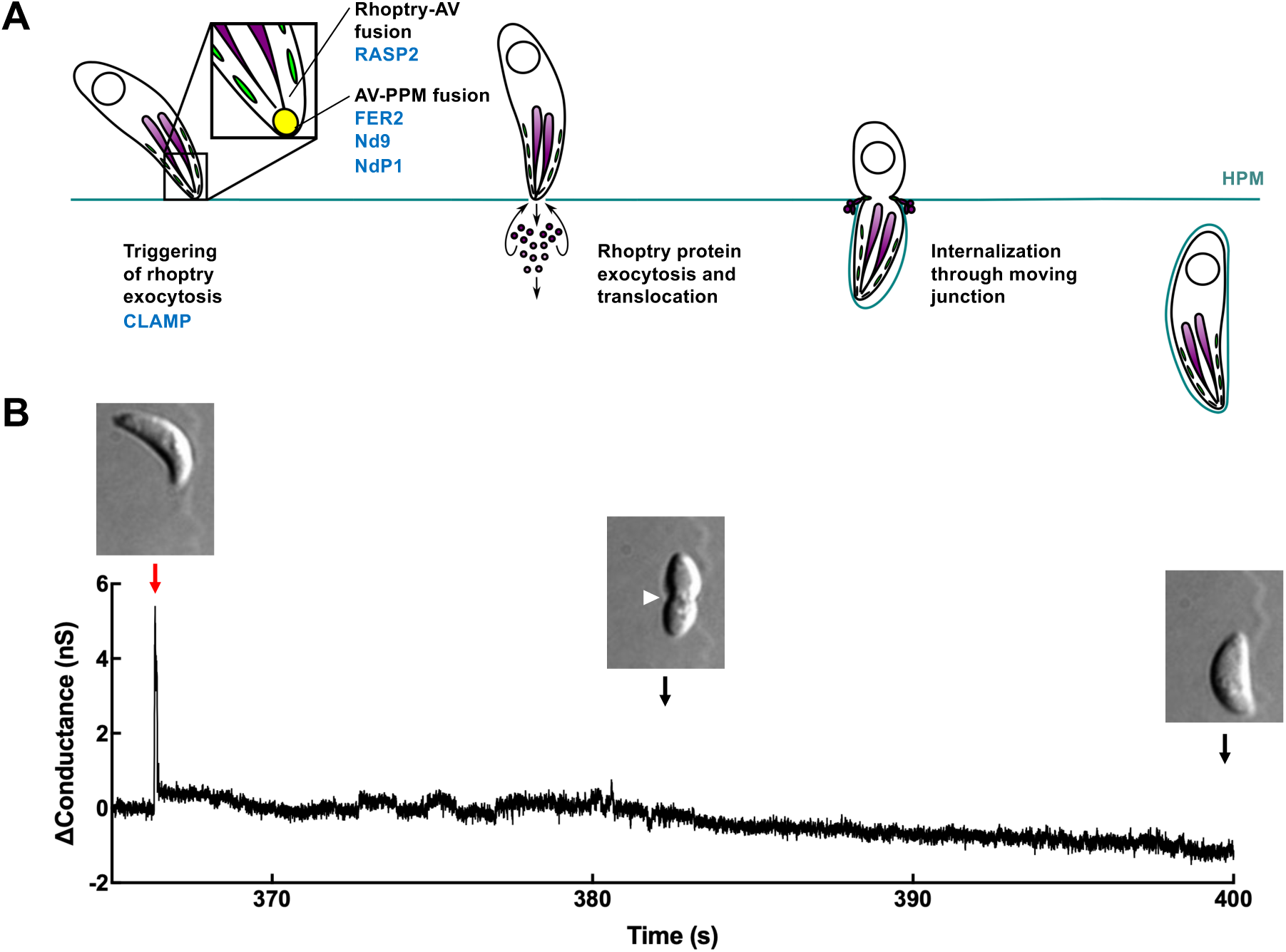
*T. gondii* invasion is associated with a transient increase in host cell plasma membrane conductance. **(A)** Schematic showing the steps of host cell invasion by *T. gondii*, highlighting rhoptry protein exocytosis and protein translocation into the host cell. The rhoptries (purple), micronemes (green), apical vesicle (yellow), and nucleus (open circle) are depicted. Specific proteins that are known to be involved in rhoptry exocytosis and are evaluated in this study are shown in blue. AV: apical vesicle; PPM: parasite plasma membrane; HPM: host plasma membrane. **(B)** Current across a COS-1 cell membrane was measured under voltage clamp conditions prior to, during, and after invasion by *T. gondii*. A single large change in host cell conductance (red arrow) is invariably observed immediately before parasite internalization, which is visualized by DIC microscopy as a constriction in the parasite plasma membrane (white arrowhead) as the parasite passes through the moving junction and into the host cell.

Despite these advances in our understanding of the composition and ultrastructure of the RSA, the mechanism by which the exocytosed rhoptry proteins are translocated into the host cell remains unknown. The underlying mechanism is likely unique to *T. gondii* and related apicomplexans. Other parasites deliver cargo into target cells via exosomes, often at a distance (Montaner et al., 2014), but there is no evidence that such a process occurs in *T. gondii*; rather, the apical localization of the RSA suggests a protein transfer mechanism that requires apposition between the parasite apex and the host cell. Prokaryotic needle-like type III, IV, and VI secretion systems are an alternative mechanism for protein delivery between cells (reviewed in Filloux, 2022; Green & Mecsas, 2016), but *T. gondii* does not possess orthologous gene products, nor have orthologous structures been observed in the many published electron micrographs of parasites prior to or during invasion (*e.g.*, Aquilini et al., 2021; Bisson et al., 2021; Dubremetz, 2007; Dubremetz, Rodriguez, & Ferreira, 1985; Gui et al., 2022; Hakansson, Charron, & Sibley, 2001; Mageswaran et al., 2021; Martinez et al., 2022; Nichols, Chiappino, & O’Connor, 1983; Sadak, Taghy, Fortier, & Dubremetz, 1988; Segev-Zarko et al., 2022; Sun et al., 2024). Fusion of the parasite and host cell plasma membranes can also be ruled out, since electrophysiology experiments have shown that host cell plasma membrane capacitance – which is a proxy for membrane surface area – does not increase during invasion (Suss-Toby, Zimmerberg, & Ward, 1996) as would be expected if the two membranes were to fuse.

A fortuitous observation made during the electrophysiology experiments referred to above provides a possible clue as to the mechanism underlying rhoptry protein translocation. The calculations of capacitance required continuous monitoring of host plasma membrane conductance, which showed an unexpected transient increase at the initiation of *T. gondii* invasion (Suss-Toby et al., 1996). We confirm this observation here and hypothesize that the change in conductance reflects a transient disruption in the barrier integrity of the host cell plasma membrane, induced by the parasite at the initiation of invasion to enable delivery of rhoptry proteins into the host cell. In an accompanying paper (Kegawa and Male et al., 2024), the electrophysiological characteristics of the invasion-associated conductance transients are analyzed in detail to explore the nature of the host cell perforation. Here, we developed an independent, fluorescence microscopy-based assay for visualizing the flux of extracellular calcium ions into the host cell through the perforation at the onset of parasite invasion. This assay was then used to test a set of mutant parasites lacking various proteins that function in rhoptry exocytosis (Fig. 1A) for their ability to induce the perforation. The results are consistent with a model in which material stored within the rhoptries is exocytosed during the initial interaction of the parasite with the host cell, causing a transient perforation in the host cell membrane through which rhoptry effector proteins are delivered.

## Results

### Host cell invasion by *T. gondii* is associated with a large, transient increase in host cell plasma membrane conductance

To confirm that the host cell plasma membrane exhibits a transient change in conductance during *T. gondii* invasion (Suss-Toby et al., 1996), COS-1 cells were patch clamped in the whole-cell configuration and held under voltage-clamp conditions (−60 mV) to measure current flowing across the host plasma membrane before, during, and after invasion. Invasion was simultaneously visualized using DIC imaging. As previously reported (Suss-Toby et al., 1996), a transient increase in host cell membrane conductance was invariably observed early in the invasion process, before moving junction formation becomes apparent as a constriction in the body of the parasite (*e.g.,* Fig. 1B). While it was important to confirm this observation, the patch clamp technique is technically demanding and its throughput is low. We therefore developed an alternative, higher throughput assay to visualize the perforation event.

### Host cell perforation can be visualized as a rapid influx of extracellular calcium into the host cell at the point of apical parasite attachment

The patch clamp method detects total current through the perforation, a physical measurement. We hypothesized that the perforation should also create a detectable chemical signal, *i.e.*, the influx of a single cation, calcium, down its steep (10,000- to 20,000-fold) concentration gradient through the perforation and into the host cell. Using the intracellular fluorescent calcium indicator, Fluo-4 AM, a transient increase in Fluo-4 fluorescence was indeed observed within the host cell at the site of invasion (Fig. 2A, top row). To unambiguously identify invading parasites, the parasites were pre-labelled with a fluorescently conjugated antibody against the most abundant surface protein, surface antigen 1 (SAG1); as the parasite invades, the bound antibody is stripped from the parasite surface at the moving junction (Dubremetz et al., 1985 and Fig. 2A, middle row). Rapid excitation switching was used for near-simultaneous imaging of the signals from the calcium indicator and the labelled parasites, enabling direct correlation of the calcium transients with specific invading parasites (Fig. 2A, bottom row and Supp. Video 1). The calcium transients within the host cell initiate at the point of apical parasite attachment, suggesting a highly localized perforation event (Fig. 2A, Supp. Video 1). Quantification of the invasion-associated calcium transient visualized in Fig. 2A is shown in Fig. 2B. While there was some variability between individual calcium transients (Supp. Fig. 1), aligning the transients by their maximal intensity reveals that the overall magnitude and kinetics of the calcium transients were consistent across independent invasion events (*e.g.*, Fig. 2C, n = 23).

**Figure 2.**
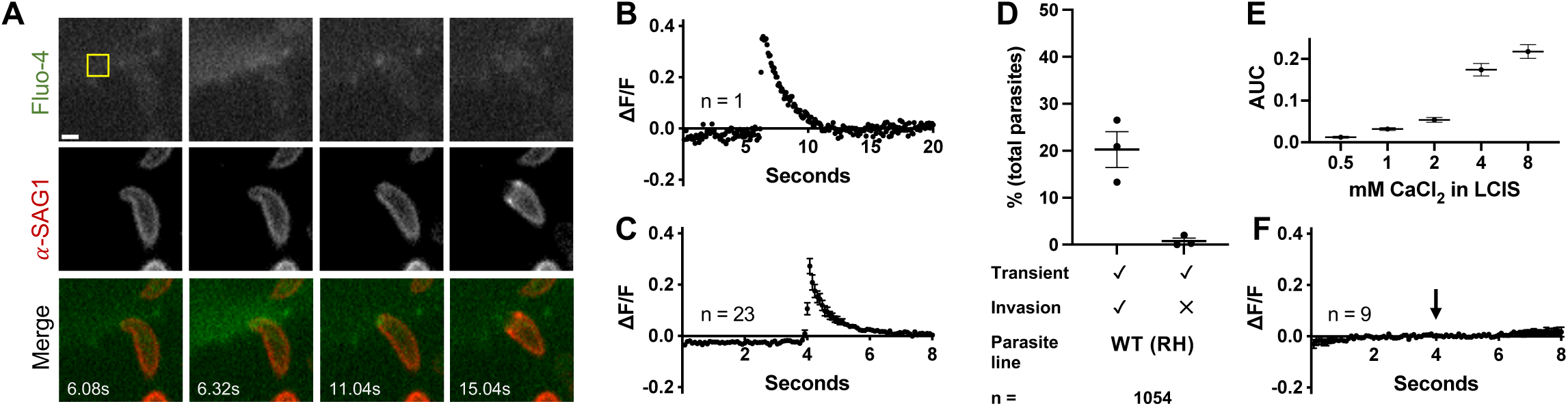
A transient increase in intracellular calcium is observed within the host cell at the site of *T. gondii* invasion. **(A)** Individual frames from a time series showing changes in Fluo-4 fluorescence and stripping of fluorescently-conjugated anti-SAG1 antibody from the surface of invading *T. gondii*. Top panels: Fluo-4 (pseudo colored green in merge); middle panels: anti-SAG1 (pseudo colored red in merge); bottom: merge. Scale bar = 2 µm. The calcium transient reaches maximal intensity at 6.32 seconds. The full video from which these frames were extracted is presented as Supp. Video 1. **(B)** Quantification of Fluo-4 fluorescence intensity in the host cell during the invasion event depicted in (A) expressed as the change in fluorescence intensity within an invasion-adjacent ROI (10×10 pixel yellow square in (A)) divided by the median intensity within the ROI over the entire time series (ΔF/F). **(C)** Consensus plot of Fluo-4 fluorescence intensities from all invasions captured in a single day (n = 23, 4 technical replicates). The fluorescence intensities in the 100 frames surrounding the peak of each calcium transient were averaged across all transients, the peaks of which were aligned to frame 51. The plot shows the mean ± SEM at each time point. **(D)** Quantification of calcium transients induced by WT (RH) parasites and whether they are associated with invasion events. Each data point represents one biological replicate consisting of the average of two to three technical replicates. Total parasite counts are shown in Supp. Figure 3A; horizontal bars indicate mean ± SEM (n = 1054 parasites). **(E)** The area under the curve (AUC) was measured for calcium transients generated in different extracellular calcium concentrations. At least 20 transients from two to three biological replicates were measured at each calcium concentration; mean ± SEM. **(F)** Consensus plot of Fluo-4 fluorescence levels from invasions in calcium-free conditions (n = 9). 100 frames were plotted for each invasion event; traces were aligned such that invasion began at 4s (arrow); mean ± SEM.

The Fluo-4 fluorescence intensity data shown in Figs. 2B and 2C were manually extracted from the video sequences, one event at a time. To improve throughput, we developed a semi-automated analysis pipeline that identifies calcium transients in an entire field of view (containing, on average, 165 ± 11 parasites) and enables subsequent visual comparison of the transients to the captured images of invading parasites (see Supp. Fig. 2, and Methods for details). Using these methods, we observed that 21.0 ± 4.4% of the wildtype (WT) RH parasites added to a host cell monolayer induced calcium transients during the 96 seconds of recording time (Fig. 2D). Of those parasites that induced transients, 98.6% subsequently invaded (Supp. Fig 3A). Large numbers of parasites can be analyzed using these methods (n = 1054 in Fig. 2D), enabling robust statistical comparisons between populations of parasites (Supp. Fig. 3).

To confirm that the calcium transients were due to influx of extracellular calcium rather than signal-mediated release of calcium from host cell intracellular stores, we conducted experiments with different concentrations of extracellular calcium. The magnitudes of the invasion-associated calcium transients were proportional to the concentration of calcium in the extracellular medium (Fig. 2E). Furthermore, during invasion in calcium-free medium no transients were detected, even though the parasites remained capable of invasion (Fig. 2F). Together, these data demonstrate that (a) the intracellular calcium transients reflect entry of extracellular calcium, and (b) the calcium transients serve as a convenient way to monitor host cell perforation, although calcium influx is not itself required for invasion.

To further validate the conductance and calcium transients as manifestations of the same perforation event, *i.e.*, to test the hypothesis that the ion flux detected via patch clamp recording occurs through the same pathway as the calcium influx detected using Fluo-4, we compared the two datasets directly. First, we generated consensus transients for conductance (Fig. 3A, n = 25) and calcium (Fig. 3B, n = 30), each denoised and peak normalized. Though on different time scales, the shapes of the transients were broadly similar, with a fast rise and slower decay. The difference in time scales of the two consensus transients reflects the specificity with which the patch clamp measurements detect changes in host plasma membrane integrity, while the calcium indicator signal is more complex, particularly in the decay phase that includes diffusion of internalized calcium and calcium-bound indicator away from the site of perforation. Since ion flux scales with the magnitude of the permeability pathway and its duration, we used the area under the curve (AUC) values of the mean normalized conductance and calcium transients to compare their distributions (Fig. 3C). There is no significant difference between these two distributions, based on a 2-sample Kolmogorov-Smirnov test (*p* = 0.73; see Methods for details).

**Figure 3.**
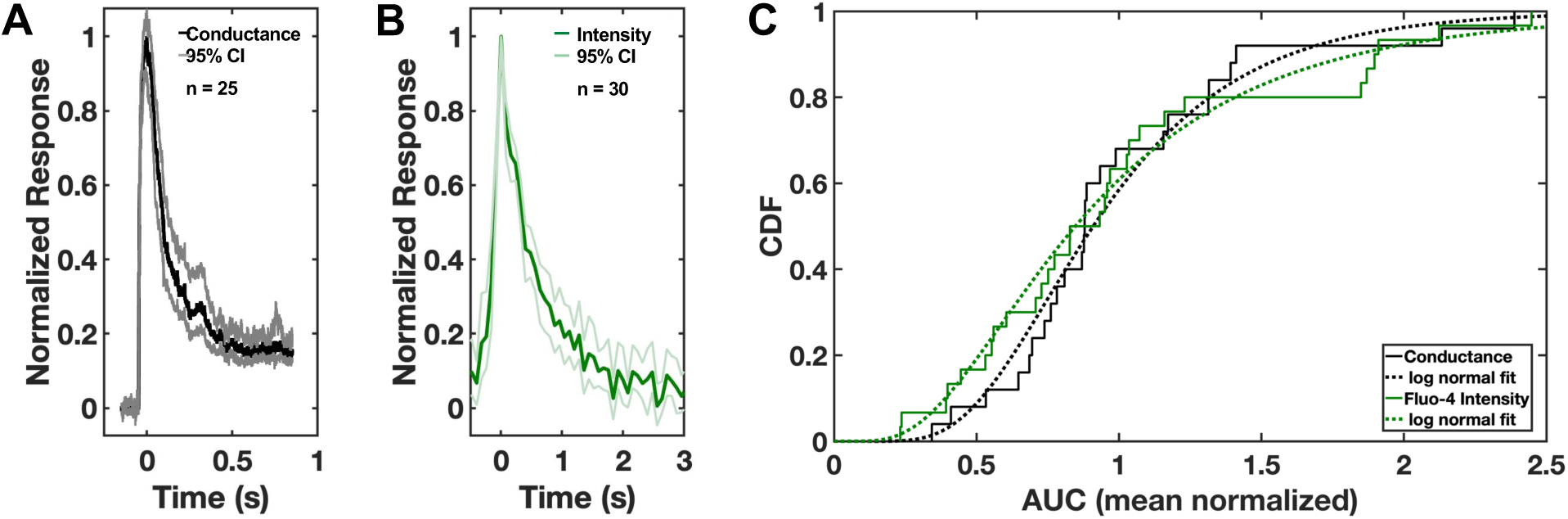
Comparison of the conductance and calcium transients induced in the host cell during invasion. **(A, B)** Consensus transients for (A) conductance (black) with 95% confidence interval (CI, grey; n = 25) and (B) Fluo-4 intensity (green) with 95% CI (light green; n = 30). The peak of each transient was normalized to 1.0 for comparison. **(C)** Area under the curve (AUC) data for the conductance (black) and Fluo-4 intensity (green) mean normalized transients were plotted as their cumulative distribution functions (CDFs), each with a log normal fit (dashed lines). *p* = 0.73, 2-sample Kolmogorov-Smirnov test.

Taken together, these results validate visualization of calcium influx as a useful alternative approach for monitoring perforation of the host cell plasma membrane during invasion. Next, we used this method and a collection of parasites that can be conditionally depleted of key proteins involved in the different steps of rhoptry exocytosis (Fig. 1A) to test whether rhoptry exocytosis is necessary for host cell perforation during invasion.

### Parasites depleted of CLAMP, which triggers rhoptry discharge in response to host cell binding, generate fewer calcium transients than wild-type parasites

CLAMP (claudin-like apicomplexan microneme protein) is a transmembrane protein involved in generating the signals that mediate rhoptry discharge, likely via recognition of as-yet unidentified molecules on the host cell surface (Sidik et al., 2016; Valleau et al., 2023). Parasites in which CLAMP is conditionally depleted by treatment with rapamycin were defective in both invasion and rhoptry protein transfer into the host cell (Sidik et al., 2016; Valleau et al., 2023). Note that the assays currently available to study rhoptry secretion are end-point assays that measure the presence of rhoptry proteins in the host cell; as such, they do not distinguish between rhoptry exocytosis and transfer of the exocytosed proteins into the host cell. We confirmed rapamycin-dependent CLAMP depletion in these parasites by Western blot (Supp. Fig. 4A). CLAMP-depleted parasites generated fewer calcium transients in the host cell compared to parasites expressing CLAMP (Fig. 4A). When the data for multiple independent biological replicates were quantified, these differences were found to be statistically significant: CLAMP-expressing parasites induced calcium transients at a rate of 22.5 ± 2.7%, compared to 4.2 ± 0.5% for CLAMP-depleted parasites (Fig. 4B; Supp. Fig. 3B, B’; *p* < 0.0001). In contrast to the CLAMP conditional knockdown parasites, rapamycin treatment had no effect on the number of calcium transients generated by the RH DiCre parental (Andenmatten et al., 2013; Pieperhoff et al., 2015) parasite line (see Methods, Supp. Fig. 5). The 4.1 ± 0.5% residual invasion and associated transient generation seen in the CLAMP-depleted parasites is consistent with the previously reported large but incomplete block in invasion associated with CLAMP depletion (Sidik et al., 2016). Thus, the loss of CLAMP leads to a reduced ability of the parasite to perforate the host cell.

**Figure 4.**
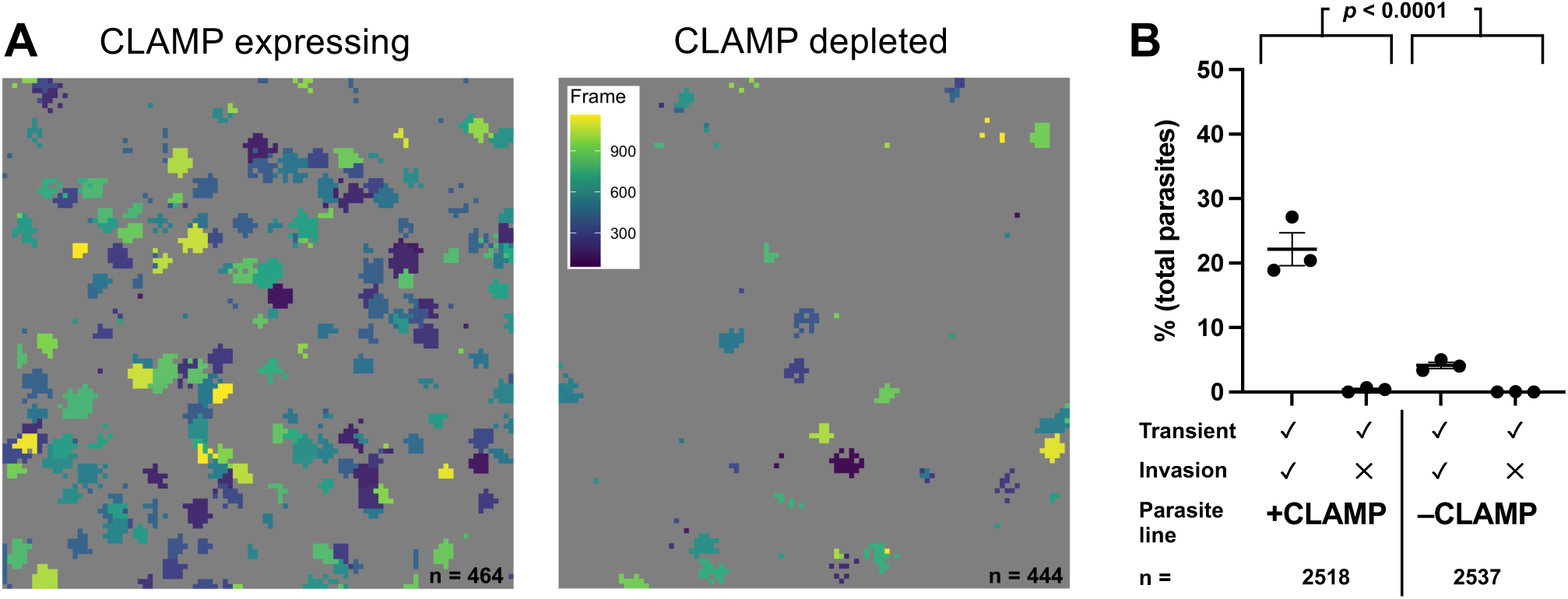
Parasites depleted of CLAMP generate fewer calcium transients than wild-type parasites. **(A)** Representative fields of view (FOV) showing calcium transients generated by CLAMP-expressing (left) and CLAMP-depleted (right) parasites (n = 464 and 444 parasites, respectively). Events are color-coded by frame of peak transient intensity (inset, right plot); FOV = 221×221 µm. **(B)** Quantification of invasion events and calcium transients induced by control parasites (+CLAMP, n = 2518) compared to parasites depleted of CLAMP (-CLAMP, n = 2537); horizontal bars indicate mean ± SEM. Each data point represents one biological replicate consisting of the average of three technical replicates. The total number of calcium transients generated by the +CLAMP and -CLAMP groups were compared using Fisher’s exact test, *p* < 0.0001 (Supp. Fig. 3B’).

### Parasites require an intact rhoptry secretory apparatus to perforate the host cell

Next, we focused on three members of the rhoptry secretory apparatus (RSA) – ferlin 2 (FER2), and two non-discharge proteins (Nd9 and NdP1) – each of which is necessary for membrane fusion between the AV and the parasite plasma membrane (Aquilini et al., 2021; Coleman et al., 2018). Protein depletion following treatment with anhydrotetracycline (ATc) was confirmed by Western blot (Supp. Fig. 4B-D). Parasites depleted of these three proteins all showed a phenotype similar to that of the CLAMP knockdown: a statistically significant reduction in host cell perforation and invasion compared to their respective controls (Fig. 5A-C; Supp. Fig. 3C-E, C’-E’; *p* < 0.0001 for each of the three lines, comparing total number of calcium transients generated by protein-depleted *vs.* control groups). The effect of ATc treatment on calcium transient generation by the RH TATi parental (Meissner, Brecht, Bujard, & Soldati, 2001) parasites was taken into consideration in these analyses (see Methods and Supp. Fig. 5). Thus, parasites depleted of proteins required for fusion of the AV and parasite plasma membrane are deficient in their ability to induce perforation of the host cell plasma membrane.

**Figure 5.**
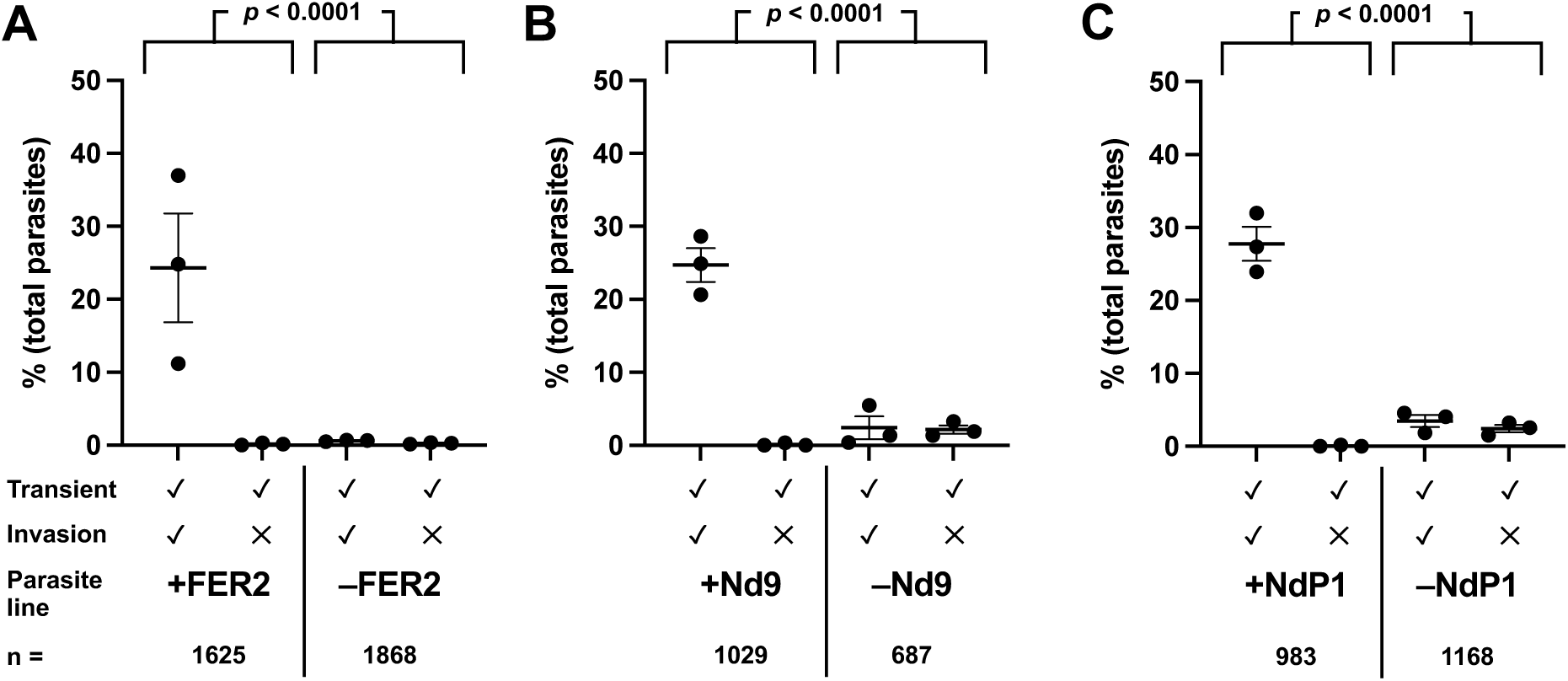
Parasites with a disrupted rhoptry secretion apparatus generate fewer calcium transients than wild-type parasites. **(A-C)** Quantitative comparison of invasion events and calcium transients induced by mutants depleted of RSA proteins involved in the AV-PPM fusion event (FER2 (A), Nd9 (B), and NdP1 (C)) and their respective controls. Each data point represents one biological replicate, consisting of the average of three technical replicates; horizontal bars indicate mean ± SEM. Total number (n) of parasites analyzed per group is indicated. The total number of calcium transients generated by the +protein and -protein groups were compared by Fisher’s exact test; *p* < 0.0001 for all the protein depleted lines (Supp. Fig. 3C’-E’).

We made two observations during these experiments that were analyzed further. First, in some of our early experiments, we noted invasions without associated calcium transients (Supp. Fig. 3). This observation seemingly argues against the hypothesis that host cell perforation is required for invasion and is distinctly different from the invariable association of the conductance transients with invasion seen by the highly sensitive patch clamp method (Kegawa and Male et al., 2024; Suss-Toby et al., 1996). To test whether calcium transients were present in these experiments but below the limit of detection of our assay, we developed an alternative assay method with improved sensitivity using a different fluorescent calcium indicator (Cal-520 AM), treatment with probenecid to improve indicator retention, and elevated levels of extracellular calcium to improve signal-to-noise. Retrospective retesting of the CLAMP inducible knockdown parasites using this improved method recapitulated the results attained with the standard method (*i.e.*, CLAMP-deficient parasites generate fewer transients; Supp. Fig. 6A, B), and, importantly, the improved method detected a reduced number of invasions without transients (0.04%; 95% confidence interval (CI, Wilson score interval) 0.01-0.21%, Supp. Fig. 6B) compared to the standard method (1.35%; 95% CI (Wilson score interval) 1.06-1.70%,, Supp. Fig. 3B). These results, together with the current recordings, suggest that few if any parasites invade without first perforating the host cell.

The second observation was that parasites occasionally generate aberrant calcium transients in the host cell. Transients induced by NdP1-depleted parasites are shown in Supp. Fig. 7A as a representative example. A small percentage of the parasites depleted of NdP1 were able to invade, as expected (Aquilini et al., 2021), and these parasites induced transients similar to those generated by WT parasites (*e.g.*, Supp. Fig. 7A, B, box i). In contrast, two parasites in the same field of view induced aberrant calcium transients (Supp. Fig. 7A, boxes ii and iii), and these parasites failed to subsequently invade. The aberrant transients had a different shape and magnitude and were therefore not well captured by our automatic peak detection software, which was originally optimized to maximize peak capture and to minimize noise of the transients induced by WT parasites. By altering the peak detection parameters, WT-like transients could still be detected (*e.g.,* Supp. Fig. 7B) and the aberrant transients (*e.g.,* Supp. Fig. 7E, G) were captured at higher frequency. The transients associated with invading NdP1-depleted parasites were similar to those generated by WT parasites (compare Supp. Fig. 7D to Fig. 2C), while the aberrant transients associated with the non-invading NdP1-depleted parasites were broader, occupying ∼10x as many pixels and showing peak fluorescence intensities approximately twice as large (Supp. Fig. 7E, F and 7G, H; Supp. Video 2) as the transients associated with invasions (Supp. Fig. 7C). These characteristics were consistent across the aberrant transients detected (Supp. Fig. 7I). Using the altered peak detection parameters, the aberrant transients were observed at a low frequency in all parasite lines tested (Supp. Fig. 7J) but were most common with Nd9- and NdP1-depleted parasites (2.1 ± 0.6 and 2.4 ± 0.5% of the parasites, respectively). The parasites that generated the aberrant transients failed to invade in >93% of the cases monitored (Supp. Fig. 7J), suggesting that the aberrant transients reflect perforations that are insufficient to support invasion.

### Parasites depleted of RASP2, which likely functions in rhoptry-to-AV fusion, are impaired in their ability to generate calcium transients

If the perforation functions in rhoptry protein translocation, the AV would be an ideal compartment in which to sequester the perforating agent, since it could be exocytosed before the bulk of the rhoptry proteins, creating the pathway for translocation across the host cell membrane immediately before the pathway is required. Given that RASP2 (rhoptry apical surface protein 2) is thought to mediate fusion between the rhoptry tip and the AV (Suarez et al., 2019), fusion of the AV to the parasite plasma membrane (and release of the perforating agent hypothetically stored within the AV) might still occur in RASP2-depleted parasites. RASP2 knockdown in the parasite line used in these experiments was confirmed by Western blot (Supp. Fig. 4E). The RASP2-depleted parasites behaved like those lacking CLAMP or members of the RSA, showing a strong reduction in the number of calcium transients induced (3.1 ± 1.3% compared to 26.4 ± 9.1% for control parasites; Fig. 6A, Supp. Fig. 3F and 3F’; *p* < 0.0001). By electrical recording, RASP2-depleted parasites failed to induce any detectable conductance transients while RASP2-expressing parasites induced conductance transients at WT levels (28.8%, 95% confidence interval 20.6 - 38.2%; Fig. 6B).

**Figure 6.**
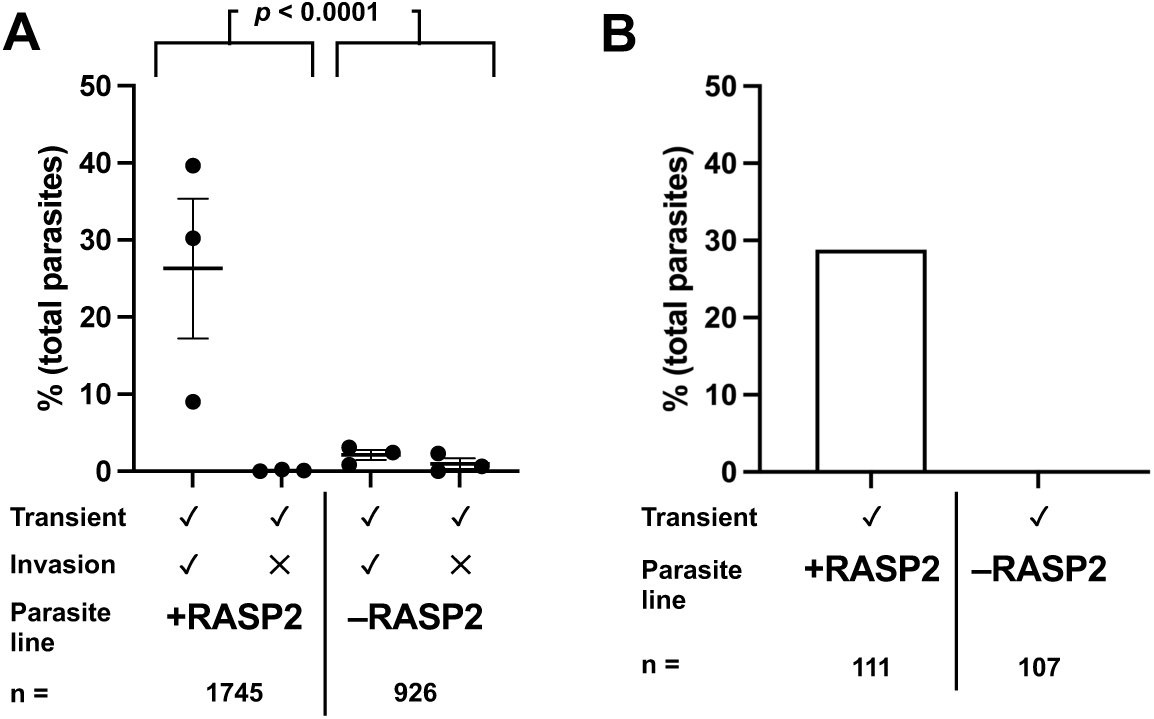
Parasites depleted of RASP2 generate fewer calcium transients than wild-type parasites. **(A)** Quantification of invasion events and calcium transients induced by control parasites (+RASP2, n = 1745) compared to parasites depleted of RASP2 (-RASP2, n = 926). Each data point represents one biological replicate, consisting of the average of three technical replicates; horizontal bars indicate mean ± SEM. The total number of calcium transients generated by the +RASP2 and -RASP2 groups were compared by Fisher’s exact test, *p* < 0.0001 (Supp. Figure 3F’). **(B)** Quantification of the number of conductance transients induced by control parasites (+RASP2, n = 111) compared to parasites depleted of RASP2 (-RASP2, n = 107), as detected by patch clamp.

## Discussion

We have shown using two independent assays that the barrier integrity of the host cell membrane is transiently disrupted during the early stages of invasion by *T. gondii*. Given the tight association between the membrane perforation and successful invasion, perforation likely plays an important role in the invasion process. We propose that the perforation provides the pathway through which exocytosed rhoptry effector proteins – including the RON proteins that are critical for moving junction formation and invasion – are delivered into the host cell. Using our new fluorescence microscopy-based assay and a collection of mutant parasites with reduced expression of proteins involved in the signaling pathway leading to rhoptry exocytosis (CLAMP), fusion of the AV to the parasite plasma membrane (FER2, Nd9, NdP1), and AV-to-rhoptry fusion (RASP2), we show that rhoptry exocytosis is necessary to generate the perforation. These data suggest that the perforating agent(s) is stored within the rhoptries and/or the AV and is therefore released only upon contact with the host cell, providing a mechanism for the parasite to sequester this potentially harmful material and release it precisely when and where it is needed (Fig. 7).

**Figure 7.**
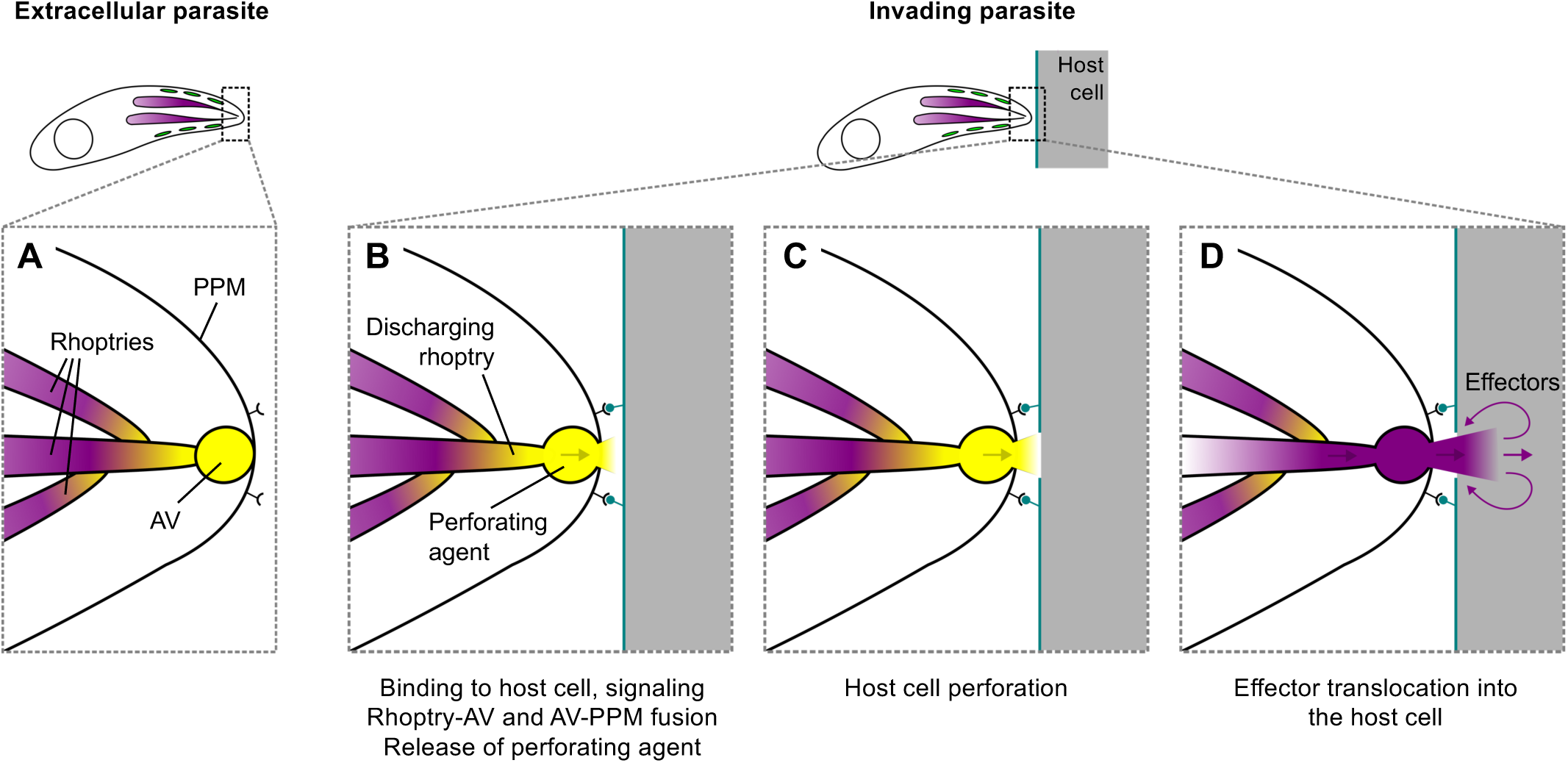
Model of rhoptry exocytosis-dependent host cell perforation. **(A)** Prior to interaction with the host cell, the perforating agent (yellow) is stored in the AV, the tip of the rhoptries, or both. **(B)** Binding of parasite surface ligands to receptor(s) on the host cell triggers signaling that leads to rhoptry exocytosis, releasing the perforating agent into the narrow space between the apical tip of the parasite and the host cell. The order in which rhoptry-to-AV and AV-to-PPM fusion take place and whether the two fusion events are independent or co-regulated are unknown. **(C)** A highly localized and transient perforation is created in the host cell membrane. **(D)** Exocytosed rhoptry effector proteins (purple) emerge from the tip of the parasite immediately after the perforating agent, and pass through the perforation and into the host cell.

The electrical and optical assays for monitoring the perforation are complementary, each with its own unique strengths. Patch clamping provides the most direct way to visualize the perforation, is highly sensitive, and offers excellent (sub-millisecond) time resolution that proved critical to our ability to resolve the transient into discrete unitary conductance events (Kegawa and Male et al., 2024). The calcium indicator assay is less arduous, provides spatial information on the location of the perforation within individual cells, and is higher throughput. The higher throughput of the fluorescence-based assay was critical to our ability to assay multiple mutant parasite lines in sufficiently large numbers (500 - 2500 parasites per treatment group) for robust statistical comparisons between parasite populations. The sensitivity of the calcium assay can be further improved for future studies using other calcium indicators (e.g. Cal-520 AM), probenecid to reduce indicator efflux, and (where appropriate) increased extracellular calcium (see Supp Fig. 6). While transient perforation of the host cell membrane appears to be important for invasion, and calcium influx can serve as a useful experimental readout for monitoring the perforating event, the influx of calcium through the perforation is not itself necessary for invasion (see Fig. 2F).

Great strides in our understanding of rhoptry exocytosis have been made in recent years through a comparison of the proteins that regulate rhoptry exocytosis in apicomplexan parasites to those that regulate trichocyst/mucocyst discharge in ciliates (reviewed in Sparvoli & Lebrun, 2021). Ciliates express many homologs of apicomplexan RSA proteins, including FER2, Nd9 and NdP1, and in both lineages depletion of these proteins leads to defective exocytosis. Similarities and differences in the ultrastructure of the secretory apparatus in the two lineages have also been informative (Aquilini et al., 2021; Mageswaran et al., 2021). One notable difference is the presence in apicomplexans of the AV between the tip of the rhoptry and the parasite plasma membrane: in the two ciliate models (*Paramecium* and *Tetrahymena*) the secretory organelles are docked directly to the plasma membrane. Rhoptry exocytosis in apicomplexans therefore requires two fusion events: rhoptry-to-AV and AV-to-parasite plasma membrane. Why might apicomplexans have evolved this extra elaboration, when the exocytotic machinery is otherwise so well conserved? Because the material exocytosed by apicomplexans (but not ciliates) is translocated into another cell, an attractive hypothesis would be that the perforating agent that functions in protein translocation is stored in the AV, as this would provide a mechanism for the perforating agent to be delivered to the host cell membrane *before* the effector proteins, creating the pathway for their subsequent translocation. In this context, the RASP2 mutant was of particular interest to test in the perforation assay, since RASP2 is thought to mediate rhoptry-to-AV fusion (Suarez et al., 2019) and AV-to-plasma membrane fusion may still occur. However, as with the RSA and signaling mutants, depleting RASP2 disrupted the parasite’s ability to perforate the host cell. The simplest explanation for this observation is that the perforating agent is stored within the rhoptries rather than the AV. Alternatively, the two fusion events may not be independent, and blocking one may interfere with the other. Because a perforating agent could potentially damage the AV or rhoptry membrane, it is also possible that the active perforating agent is composed of two inactive components, one stored in the AV and one stored in the neck of the rhoptries. In this scenario the perforating activity would only be generated when the two compartments mix as a result of rhoptry-to-AV fusion, which in *Plasmodium* likely occurs before exocytosis of the rhoptry contents (Martinez et al., 2022). Resolution of these questions will ultimately require the identification of the perforating agent and its localization within the parasite prior to interaction with a host cell.

The mechanisms and proteins underlying host cell invasion appear to be well conserved among apicomplexan parasites. Conserved proteins include those that regulate rhoptry exocytosis, such as RASP2 and Nd9, and depletion of these proteins leads to similar phenotypes in *T. gondii* and *Plasmodium* spp. (Aquilini et al., 2021; Suarez et al., 2019). Intriguingly, a previous report demonstrated that the invasion of Fluo-4-loaded erythrocytes by *P. falciparum* merozoites is occasionally accompanied by “a strong Ca^2+^ flux spreading into the erythrocyte from the invasion site” (Weiss et al., 2015), suggesting that the membrane perforation event that we have described here also occurs during invasion by malaria parasites. In turn, we occasionally observed a phenomenon described in this previous report: a Fluo-4 signal appears at the parasite apex and is visible for a short time at the initiation of invasion (Weiss et al., 2015; Supp. Fig. 8 and Supp. Videos 1, 3-5). This observation suggests transient continuity and bidirectional transfer of material between the host cell cytosol and an interior compartment of the parasite, likely the rhoptries, and is consistent with the hypothesis that the host perforation plays a role in transfer of rhoptry effector proteins into the host cell.

We also observed another rare but intriguing phenotype where a small subset of the parasites interacting with host cells induced aberrant calcium transients that had larger amplitudes and longer durations than those typically induced by invading parasites. These aberrant calcium transients were observed most frequently in mutants depleted of proteins involved in fusion of the AV and parasite plasma membrane (Nd9, NdP1) and more rarely in the other parasite lines studied. Most of these aberrant transients (>93%) were associated with parasites that failed to subsequently invade. Why might the magnitude and kinetics of this subset of transients be different? Closure of the putative pores may require exocytosed rhoptry protein(s) and would therefore be delayed if exocytosis is disrupted or if the proteins are delivered in a less focal, concentrated bolus. Alternatively, if rhoptry effector proteins are normally translocated into the host cell through the perforation, during translocation these effector proteins will likely cause partial occlusion of the putative pore, thereby reducing calcium influx. In the case of the aberrant transients, perhaps perforation occurs without the normal levels of effector exocytosis, reducing occlusion of the putative pores and resulting in more calcium entry.

In summary, the data presented here show that rhoptry exocytosis is required for parasite-induced perforation of the host cell membrane during invasion, providing a mechanism for the parasite to release the perforating agent precisely when and where it is needed. The accompanying paper (Kegawa and Male et al., 2024) begins to address the nature of the perforating agent through a detailed electrophysiological characterization of the parasite-induced perforation event and an exploration of its mechanism. Efforts are currently underway to identify the perforating agent itself, which will enable a direct test of the hypothesis that the perforation serves as the conduit through which rhoptry effector proteins are delivered into the host cell. Given the central role played by many of the secreted rhoptry effector proteins in parasite virulence (Saeij et al., 2006; Shwab et al., 2016; Taylor et al., 2006), elucidating the mechanism(s) underlying effector entry is not only of fundamental cell biological interest, but may also identify new targets and inspire new strategies for therapeutic development. Rather than targeting a single translocated rhoptry protein, targeting the rhoptry protein delivery mechanism will simultaneously disrupt transfer into the host cell of many of the effector proteins that play such a central role in the pathogenesis of the devastating diseases caused by *T. gondii* and other apicomplexan parasites.

## Methods

### Parasite and cell culture

*T. gondii* parasite lines were propagated by serial passage in confluent monolayers of human foreskin fibroblast (HFF) cells. HFFs were maintained in Dulbecco’s Modified Eagle Medium (DMEM) (Life Technologies, Carlsbad, CA) with 10% v/v heat-inactivated fetal bovine serum (FBS) (Life Technologies, Carlsbad, CA), 10 mM HEPES pH 7.0, 100 units/mL penicillin, and 100 µg/mL streptomycin. Prior to parasite passage, the medium was replaced with DMEM with 1% v/v FBS, 10 mM HEPES pH 7.0, 100 units/mL penicillin, and 100 μg/mL streptomycin.

DiCre/CLAMP parasites were treated with 50 nM rapamycin (Invitrogen, Waltham, MA) to induce CLAMP depletion or an equivalent volume of DMSO for 2 hours prior to 48-hour culture in drug-free medium (Sidik et al., 2016). All other inducible knockdown (iKD) parasite lines were pretreated with 1.5 µg/mL anhydrotetracycline (ATc) (Takara Bio, San Jose, CA) to induce gene knockdown or an equivalent volume of 100% ethanol (EtOH) for the following time courses prior to experiments: FER2 iKD, 96 hours (Coleman et al., 2018); Nd9 iKD, 72 hours (Aquilini et al., 2021); NdP1 iKD, 72 hours (Aquilini et al., 2021); and RASP2 iKD, 48 hours (Suarez et al., 2019). Experiments with Nd9 iKD parasites were performed in the continuous presence of 1.5 µg/mL ATc or EtOH.

To isolate parasites for experiments, DMEM containing 1% FBS was replaced with Endo buffer (modified to include calcium: 44.7 mM K_2_SO_4_, 8 mM MgSO_4_, 2 mM CaSO_4_, 106 mM sucrose, 5 mM glucose, 20 mM Tris-H_2_SO_4_, 3.5 mg/mL BSA) (Endo, Tokuda, Yagita, & Koyama, 1987) before HFFs containing large vacuoles were detached from the flask using a cell scraper and parasites released by 2-3 passages through a 26G 1/2” blunt needle. Parasites were isolated from cell debris by passage through a Whatman Nuclepore Track-Etch Membrane 3 µm filter (MilliporeSigma, Burlington, MA), and spun at 1000×g for 2 minutes. Parasites were then resuspended in 1:20 anti-SAG1 antibody (monoclonal antibody DG52, a generous gift from Dr. David Sibley, 0.2 mg/mL stock) conjugated to AlexaFluor 647 (Alexa Fluor™ 647 Antibody Labeling Kit (Molecular Probes, Eugene, OR)) in Endo buffer for 30 minutes at ambient temperature, then spun at 1000×g for 2 minutes, and resuspended in Endo buffer at 3×10^7^ parasites/mL.

### Live calcium transient / invasion assay

#### Assay preparation

HFFs were seeded in 3 chambers of an ibidi µ-Slide VI 0.4 (ibidi GmbH, Gräfelfing, Germany) to an average final density of 90% in DMEM containing 10% FBS and incubated for 2 hours at 37°C (with 5% CO_2_ and humidity). HFFs were washed 1× with live cell imaging solution (LCIS) (Ringer’s solution with glucose to 20 mM: 155 mM NaCl, 3 mM KCl, 2 mM CaCl_2_, 1 mM MgCl_2_, 3 mM NaH_2_PO_4_, 10 mM HEPES, 20 mM glucose) before being loaded with 5 µM Fluo-4 AM (Invitrogen, Waltham, MA) with 1% PowerLoad Concentrate, 100X (Invitrogen, Waltham, MA) in LCIS for 50 minutes at ambient temperature. Cells were washed 1× with LCIS to wash out excess indicator and incubated for an additional 40 minutes at ambient temperature to allow complete de-esterification of intracellular AM esters.

For Supp. Fig. 6, Supp. Video 4, and Supp. Video 5, the assays were performed as above except that HFFs were loaded with 5 µM Cal-520 AM (AAT Bioquest, Pleasanton, CA) with 1% PowerLoad Concentrate, 100X (Invitrogen, Waltham, MA), and 1 mM probenecid (Invitrogen, Waltham, MA) for 90 minutes at 37°C followed by 30 minutes at ambient temperature. All subsequent solutions included 1 mM probenecid. Cells were washed 1× with LCIS to wash out excess indicator.

#### Imaging

Experiments were carried out at 35-36°C. Fluo-4-loaded cells were washed 1× with Endo buffer before allowing pre-labeled parasites to settle on cells for 10 minutes. The buffer was then exchanged for pre-warmed invasion-permissive LCIS to capture calcium transients and invasion events.

Imaging was carried out on a Nikon Eclipse TE300 widefield epifluorescence microscope (Nikon Instruments, Melville, NY) using a 60× PlanApo λ objective (0.22 µm/pixel, NA 1.4). 1020×1020 pixel images were captured using an iXon 885 EMCCD camera (Andor Technology, Belfast, Ireland) set to trigger mode, with exposure time of 39 ms, no binning, readout speed of 30 MHz, conversion gain of 3.8×, and EM gain of 300. Perforation of host cells by parasites resulting in calcium transients and subsequent invasion events were observed by near-simultaneous excitation of Fluo-4 or Cal-520 (490 nm) and AlexaFluor 647 (635 nm) using a pE-4000 LED illumination system (CoolLED, Andover, England), through rapid excitation switching triggered by the NIS Elements Illumination Sequence module (Nikon Instruments, Melville, NY). Hardware was driven by NIS Elements v. 5.11 software (Nikon Instruments, Melville, NY).

#### Data processing

1200 high quality frames (96 seconds) were processed from each capture. Mean fluorescence intensity (MFI) was extracted from 10,404 10×10 pixel regions of interest (ROIs) over time from each 1020×1020 pixel field of view (FOV) in ImageJ (Supp. Fig. 2A; 100×100 pixel FOV containing a single calcium transient shown for clarity in Supp. Fig. 2A-C). MFI results were passed through PeakCaller script in MATLAB (MathWorks, Natick, MA) for automated identification of intracellular calcium transients (Artimovich, Jackson, Kilander, Lin, & Nestor, 2017). Using RStudio, calcium transient results from each ROI were plotted back to ROI location and color-coded, based on both the time (frame) the transient reached its peak (Supp. Fig. 2B), and the amplitude (peak) of the transient (Supp. Fig. 2C). Calcium transient and invasion events were quantified through comparison of identified peaks and live image captures (Supp. Fig. 2D, 600 of 1200 frames shown for clarity). For mutant parasite experiments, three biological replicates were carried out for each mutant. Each biological replicate consisted of three technical replicates for protein-depleted mutants and three technical replicates for controls, carried out on the same day. For the WT RH parasite experiments (Fig. 2D), one technical replicate from one biological replicate was excluded from downstream analysis due to poor quality.

#### Statistical analysis

For each mutant parasite line, differences between protein-depleted parasites compared to controls with respect to induction of calcium transients and invasions were assessed using Fisher’s exact test for contingency table analysis. The null hypothesis in each case was that control and protein-depleted parasites would show no differences in the counts observed for each category. The data from three biological replicates were combined into four categories (+transient / +invasion, +transient / –invasion, –transient / +invasion, and –transient / –invasion) and compared using 4×2 contingency tables (Supp. Fig. 3B-F).

The specific difference in the frequency of calcium transients induced by protein-depleted parasites compared to controls was assessed using Fisher’s exact test for the analysis of 2×2 contingency tables. The null hypothesis in each case was that control and protein-depleted parasites are equally likely to induce calcium transients. The data from the three biological replicates were combined and the number of transients generated (by invading and non-invading parasites combined) were compared using 2×2 contingency tables (Supp. Fig. 3B’-F’). Differences reported in the text as “statistically significant” refer to comparisons in which the null hypothesis was rejected based on the reported *p*-value.

Fisher’s exact test was also applied to 4×2 and 2×2 contingency tables in which counts were corrected based on the percent confluency of the host cell monolayer for each technical replicate. Zero values were treated conservatively (Jovanovic & Levy, 1997) (3/n adjustment). The resulting *p-*values remained consistent with the uncorrected count data (Supp. Fig. 3). Additionally, Fisher’s exact test was carried out on 4×2 and 2×2 contingency tables, both with and without host cell confluency correction, for each biological replicate individually. For all tests, *p* < 0.0001, except for one Nd9 biological replicate 2×2 host cell confluency-adjusted contingency table for which *p* = 0.0051.

### Control experiments conducted on parental cell lines

Control live calcium transient/invasion experiments were conducted on the RH DiCre parental line after rapamycin or DMSO treatment as described above. Parasites treated with rapamycin or DMSO both induced calcium transients and invaded (Supp. Fig. 5A-B). Rapamycin treatment did not have a significant effect on induction of calcium transients (*p* = 0.75, Fisher’s exact test; Odds ratio (95% CI): 1.02 (0.85, 1.22); Supp. Fig. 5B’).

Control live calcium transient/invasion were conducted on the RH TATi parental line after ATc or ethanol (EtOH) treatment as described above. Parasites treated with ATc or EtOH both induced calcium transients and invaded (Supp. Fig. 5C-D). ATc treatment was found to have a significant effect on induction of calcium transients (*p* < 0.0001, Fisher’s exact test; Odds ratio (95% CI): 0.57 (0.44, 0.73), *p* < 0.001; Supp. Fig. 5B’). To determine the effect of this result on the TATi inducible knockdown (iKD) lines, the Breslow-Day test was used to analyze the interaction between the odds ratio from the TATi control experiment to the odds ratio from each of the iKD line experiments, with the null hypothesis that the odds ratios are equal. For all comparisons, the odds ratios were found to be significantly different (*p* < 0.001, Breslow-Day test; Supp Fig. 5E). Further, by converting these odds ratios to effect sizes, the effect size for ATc treatment in the TATi dataset was found to be small-to-moderate (0.2-0.5; Supp Fig. 5E) while the effect sizes for gene knockdown within all of the iKD datasets were all found to be large (>0.5; Supp. Fig. 5E)(Chinn, 2000).

Due to these large differences in effect size, we conclude that the differences observed within each iKD line dataset and presented in the figures are largely due to gene knockdown.

### Electrophysiology experiments

COS1 cells (ATCC CRL-1650) were cultured in DMEM supplemented with high glucose, 200 mM Glutamax, 1 mM sodium pyruvate (ThermoFisher Scientific, Cat#10569, Waltham, MA) and 100 μg/ml primocin (InvivoGen Cat#ant-pm-2, San Diego, CA) with 10% FBS Premium Select (R&D Systems, Cat#S11550, Minneapolis, MN) at 37°C under 5% CO2. For experiments, COS-1 cells were seeded onto a 35 mm DT dish (Bioptechs, Butler, PA) at a concentration of 4×10^4^ cells/mL in 1 mL DMEM for 60 minutes at 37°C under 5% CO_2_. before the medium was replaced by LCIS. Temperature was maintained at 37°C using a Delta T4 Culture Dish Controller (Bioptechs, Butler, PA). The pipette solution for whole cell recordings contained 122 mM KCl, 2 mM MgCl_2_, 11 mM EGTA, 1 mM CaCl_2_, 5 mM HEPES (pH 7.26, adjusted with KOH). Patch pipettes (3 MΩ resistance) were fabricated from 1.5-mm thick wall borosilicate glass capillaries using a P1000 puller (Sutter Instruments, Novato, CA). Whole cell formation was monitored with an AxoPatch200B amplifier (Molecular Devices, San Jose, CA) in the voltage clamp mode. The output current was filtered using the internal 100 kHz Lowpass Bessel filter included in the amplifier and an external 5 kHz low pass, 8 pole, Bessel filter (Model 900 CT/9 L8L, Frequency Devices Inc, Haverhill, MA) and was digitized at 100 μs for a time resolution of 200 μs. A -60-mV holding potential was applied to monitor current changes during interactions with *T. gondii* parasites. Current was recorded for a maximum of 15 minutes per cell, digitized using an Axon Digidata 1550B (Molecular Devices, San Jose, CA) and the Axopatch software package (Molecular Devices, San Jose, CA). Conductance is calculated using Ohm’s law following correction for the pipette access resistance. Data analysis was performed offline using Clampfit 11.2 (Molecular Devices, San Jose, CA) and MatLab r2022b (MathWorks, Natick, MA).

### Comparisons between optical and electrical transients

Raw fluorescence trajectories (n = 22; 160 second duration; 50×50 pixel ROI), with variable numbers of transients were wavelet denoised (Daubechies db6 wavelet, empirical Bayes denoising threshold) and the time varying background calculated on the denoised trajectory using the rolling ball algorithm in time (disc structural element, 2 sec radius). This background was then subtracted from the raw fluorescence trajectory. In the absence of a transient, the background subtracted baseline was characterized by an approximately zero mean gaussian (0.56) indicating less than 1 intensity unit bias and consistent with appropriate background subtraction. To rapidly identify robust transients, a threshold of 5*(Standard Deviation) was used with MATLAB peakfinder; 30 transients were identified in this data set, peak normalized, and the peak arbitrarily set to zero time (Fig. 3). Conductance transients were displayed similarly. Area under the curve (AUC) was calculated by summing the background subtracted optical and conductance values from the start of the transient for a defined time (900 msec for conductance, 3 seconds for fluorescence). The distributions around the mean normalized AUC for calcium and conductance transients were compared using a 2-sample Kolmogorov-Smirnov test. Results reported in the text as not significant refer to comparisons in which there is insufficient evidence to reject the null hypothesis that the data came from populations with the same distribution.

## Acknowledgements

We thank Dr. Vern Carruthers for helpful early discussions on this project and members of the Ward lab for comments on the manuscript. This work was supported by U.S. Public Health Service grants U01AI169067 (GEW) and R01AI144369 (SL), European Research Council Advanced Grant 833309 (ML), and the intramural program of the NICHD. YK was supported by a JSPS research fellowship for Japanese Biomedical and Behavioral researchers at NIH from 2021 to 2023. The authors declare that they have no conflict of interest.

