## Supplemental figures for "Perforation of the host cell plasma membrane during *Toxoplasma gondii* invasion requires rhoptry exocytosis"

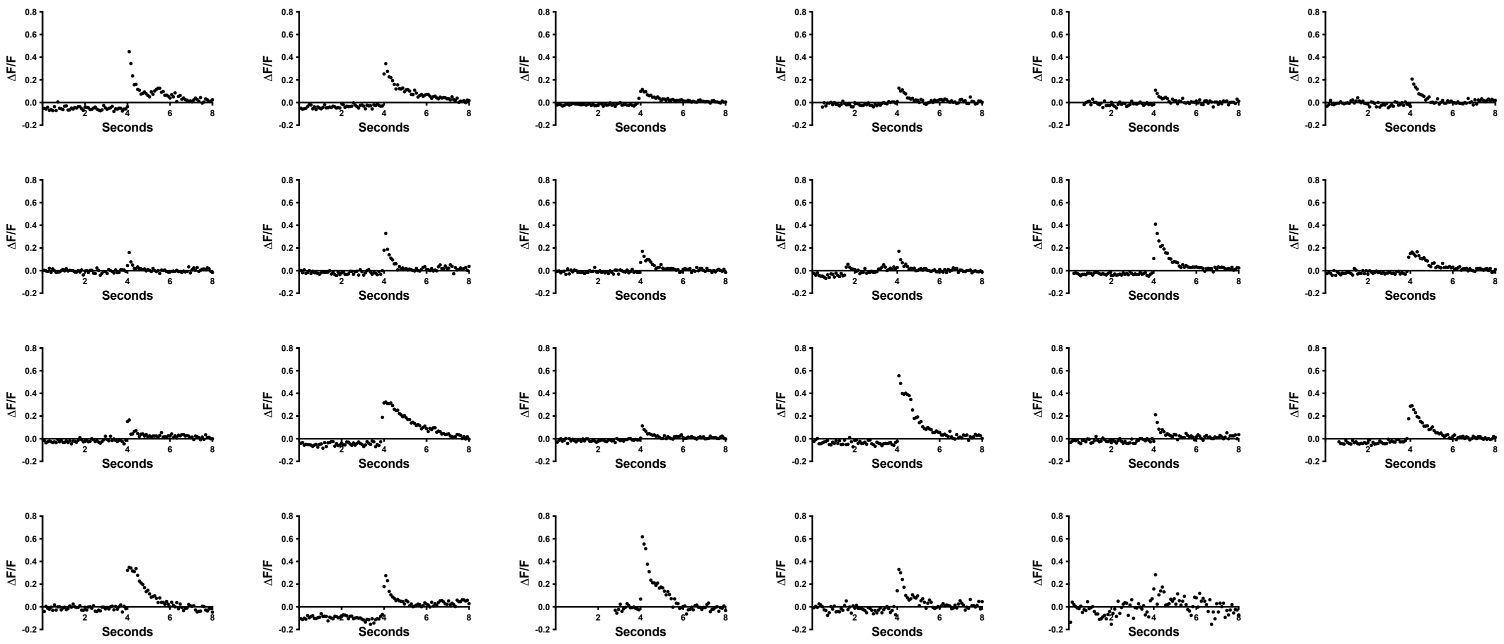


**Supp. Figure 1. Gallery of the 23 individual calcium spikes combined to generate Fig. 2C.**
Fluo-4 fluorescence intensities (ΔF/F) in the host cell measured during 23 individual *T. gondii* invasion events are shown.

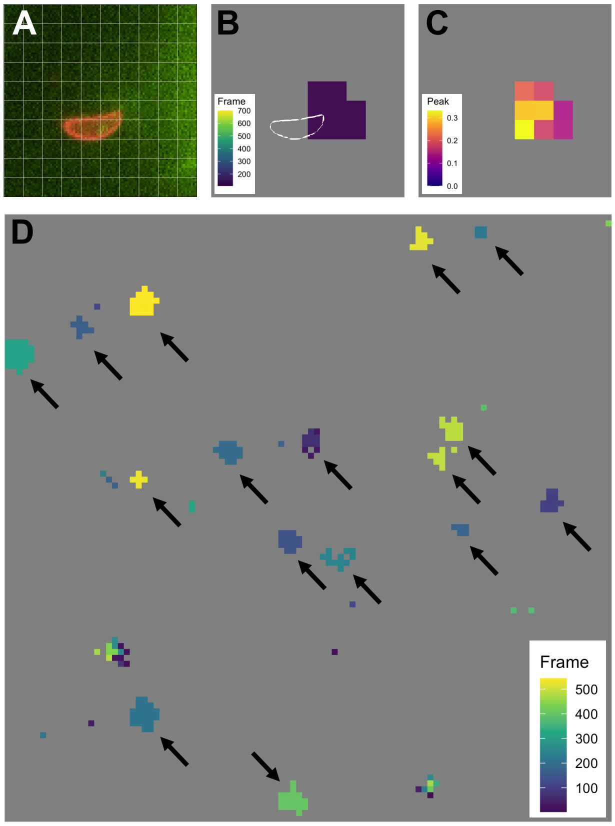


**Supp. Figure 2. Semi-automated workflow for quantifying the parasite-induced calcium transients in the host cell.**

**(A)** Individual invasion event used to illustrate the quantification scheme. Mean fluorescence intensity (MFI) was captured over time from 100 10×10 pixel regions of interest (ROIs; white grid lines) in this 100×100 pixel example field of view (FOV). The full FOV for each capture consists of 1020×1020 pixels.

**(B)** PeakCaller output plotted back to ROI location and color-coded by frame (inset) at which the transient event reached maximal intensity; location of parasite outlined in white. This transient reached its peak at frame 106 of 700 frames total.

**(C)** PeakCaller output plotted back to ROI location and color-coded by peak intensity (inset) achieved during the transient event.

**(D)** 16 calcium transients (black arrows) were detected in this representative full FOV (221×221 μm) over the course of 48 seconds of recording (600 frames). Full captures consist of 1200 frames; fewer frames are shown here for clarity. MFI was captured over time from 10,404 10×10 pixel ROIs across 1020×1020 pixel FOV. PeakCaller output was plotted back to ROI location and color-coded by the frame (inset) when each transient reached peak intensity.

**A WT (RH)**

| Treatment | +Transient +Invasion | +Transient –Invasion | –Transient +Invasion | –Transient –Invasion |
|---|---|---|---|---|
| N/A | 206 | 3 | 9 | 836 |

**B CLAMP**

| Treatment | +Transient +Invasion | +Transient –Invasion | –Transient +Invasion | –Transient –Invasion | Fisher's exact test |
|---|---|---|---|---|---|
| DMSO | 521 | 9 | 57 | 1934 | *p* < 0.0001 |
| Rapamycin | 113 | 1 | 11 | 2412 | |

**B'**

| +Transient | –Transient | Fisher's exact test |
|---|---|---|
| 530 | 1991 | *p* < 0.0001 |
| 114 | 2423 | |

**C FER2**

| Treatment | +Transient +Invasion | +Transient –Invasion | –Transient +Invasion | –Transient –Invasion | Fisher's exact test |
|---|---|---|---|---|---|
| EtOH | 436 | 3 | 21 | 1165 | *p* < 0.0001 |
| ATc | 11 | 4 | 1 | 1852 | |

**C'**

| +Transient | –Transient | Fisher's exact test |
|---|---|---|
| 439 | 1186 | *p* < 0.0001 |
| 15 | 1853 | |

**D Nd9**

| Treatment | +Transient +Invasion | +Transient –Invasion | –Transient +Invasion | –Transient –Invasion | Fisher's exact test |
|---|---|---|---|---|---|
| EtOH | 261 | 1 | 56 | 711 | *p* < 0.0001 |
| ATc | 10 | 16 | 7 | 654 | |

**D'**

| +Transient | –Transient | Fisher's exact test |
|---|---|---|
| 262 | 767 | *p* < 0.0001 |
| 26 | 661 | |

**E NdP1**

| Treatment | +Transient +Invasion | +Transient –Invasion | –Transient +Invasion | –Transient –Invasion | Fisher's exact test |
|---|---|---|---|---|---|
| EtOH | 260 | 1 | 14 | 708 | *p* < 0.0001 |
| ATc | 42 | 29 | 6 | 1091 | |

**E'**

| +Transient | –Transient | Fisher's exact test |
|---|---|---|
| 261 | 722 | *p* < 0.0001 |
| 71 | 1097 | |

**F RASP2**

| Treatment | +Transient +Invasion | +Transient –Invasion | –Transient +Invasion | –Transient –Invasion | Fisher's exact test |
|---|---|---|---|---|---|
| EtOH | 384 | 2 | 23 | 1336 | *p* < 00001 |
| ATc | 18 | 6 | 0 | 902 | |

**F'**

| +Transient | –Transient | Fisher's exact test |
|---|---|---|
| 386 | 1359 | *p* < 0.0001 |
| 24 | 902 | |

**Supp. Figure 3. Total invasion and transient counts for each parasite line analyzed and contingency table analysis for the inducible knockdown parasites.**

**(A)** Total WT parasite counts were separated into four categories: +transient / +invasion; +transient / –invasion; –transient / +invasion; and –transient / –invasion. Each number represents the sum of 3 biological replicates, consisting of 2-3 technical replicates each.

**(B-F)** Data from all mutant parasite lines categorized as in (A) within 4×2 contingency tables to compare control (top row) and protein-depleted (bottom row) parasites. Each number represents the sum of 3 biological replicates, consisting of 2-3 technical replicates each. Fisher's exact test was used for the comparison (right-most column).

**(B'-F')** Data from each mutant line (B-F) were summed based on the presence or absence of detected calcium transients (+transient, –transient) and organized into 2×2 contingency tables. Fisher's exact test was used for the comparison (right-most column).

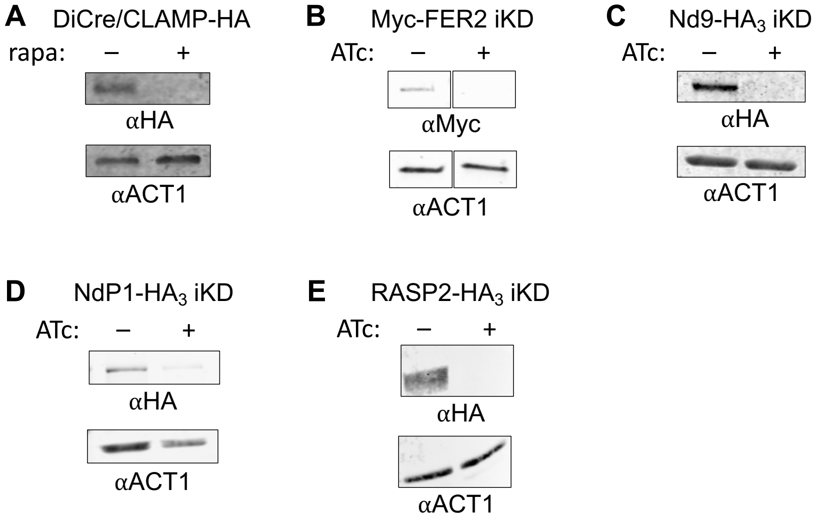


**Supp. Figure 4. Confirmation of protein depletion in the mutant parasite lines by Western blot.**

**(A)** DiCre/CLAMP-HA parasites were treated with 50 nM rapamycin (rapa; +) or an equivalent volume of DMSO (-) for 2 hours prior to 48-hour culture in drug-free medium, then total parasite lysate was analyzed by Western blot using anti-HA. Anti-actin (ACT1) was used as a loading control.

**(B-E)** Parasite lines were treated with 1.5 µg/mL anhydrotetracycline (ATc; +) or an equivalent volume of EtOH (-) for 96 hours (Myc-FER2 iKD (B)), 72 hours (Nd9-$HA_3$ iKD (C), NdP1-$HA_3$ iKD (D)), or 48 hours (RASP2-$HA_3$ iKD (E)), then total parasite lysate was analyzed by Western blot using either anti-myc or anti-HA. Anti-ACT1 was used as a loading control.

**A**

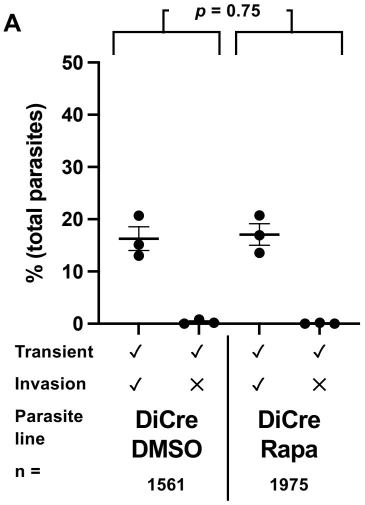


**C**

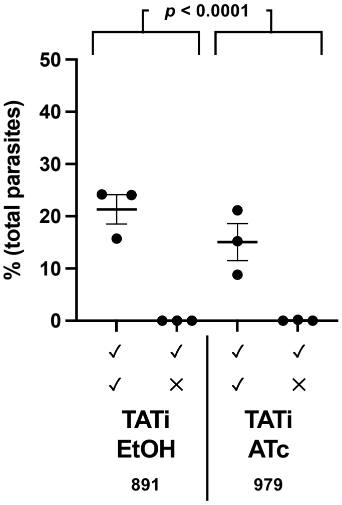


**B** **RH DiCre**

| Treatment | +Transient +Invasion | +Transient –Invasion | –Transient +Invasion | –Transient –Invasion | Fisher's exact test |
|---|---|---|---|---|---|
| DMSO | 252 | 3 | 2 | 1304 | $p$ = 0.0069 |
| Rapamycin | 330 | 1 | 17 | 1627 | |

**B'**

| Treatment | +Transient | –Transient | Fisher's exact test | Odds ratio (95% CI) | Odds ratio P-value |
|---|---|---|---|---|---|
| DMSO | 255 | 1306 | $p$ = 0.75 | 1.02 (0.85, 1.22) | $p$ = 0.838 |
| Rapamycin | 331 | 1664 | | | |

**D** **RH TATi**

| Treatment | +Transient +Invasion | +Transient –Invasion | –Transient +Invasion | –Transient –Invasion | Fisher's exact test |
|---|---|---|---|---|---|
| EtOH | 178 | 0 | 38 | 675 | $p$ < 0.0001 |
| ATc | 121 | 1 | 27 | 830 | |

**D'**

| Treatment | +Transient | –Transient | Fisher's exact test | Odds ratio (95% CI) | Odds ratio P-value |
|---|---|---|---|---|---|
| EtOH | 178 | 713 | $p$ < 0.0001 | 0.57 (0.44, 0.73) | $p$ < 0.001 |
| ATc | 122 | 857 | | | |

**E**

| Line | Odds ratio (95% CI) | Breslow-Day test (compared to TATi) | Effect size (absolute value) |
|---|---|---|---|
| TATi (Supp. Fig. 8D') | 0.57 (0.44, 0.73) | -- | 0.311 |
| FER2 (Supp. Fig. 3C') | 0.02 (0.01, 0.04) | $p$ < 0.001 | 2.161 |
| Nd9 (Supp. Fig. 3D') | 0.08 (0.05, 0.13) | $p$ < 0.001 | 1.395 |
| NdP1 (Supp. Fig. 3E') | 0.18 (0.13, 0.23) | $p$ < 0.001 | 0.947 |
| RASP2 (Supp. Fig. 3F') | 0.09 (0.06, 0.13) | $p$ < 0.001 | 1.330 |

**Supp. Figure 5. The effects of rapamycin or ATc treatment on calcium transients and invasions for parental parasite lines.**

**(A)** Quantification of invasion events and calcium transients induced by RH DiCre parental line parasites treated with DMSO (DiCre DMSO, n = 1561) compared to parasites treated with rapamycin (DiCre Rapa, n = 1975). Each data point represents one biological replicate, consisting of the average of 2-3 technical replicates; horizontal bars indicate mean ± SEM. Comparison of total calcium transients between DMSO and rapamycin groups was analyzed using Fisher's exact test, $p$ = 0.75 (see panel B').

**(B)** Total data from DMSO-treated (top row) and rapamycin-treated (bottom row) RH DiCre parasites were categorized as in Supp. Fig. 3A-F within a 4×2 contingency table for comparison. Fisher's exact test was used for the comparison (right-most column). Each number represents the sum of 3 biological replicates, consisting of 2-3 technical replicates each.

**(B')** Data from (B) were summed based on the presence or absence of detected calcium transients (+ transient, –transient) and organized into 2×2 contingency tables. Fisher's exact test and odds ratio were used for the comparison.

**(C)** Quantification of invasion events and calcium transients induced by RH TATi parental line parasites treated with ethanol (TATi EtOH, n = 891) compared to parasites treated with anhydrotetracycline (TATi ATc, n = 979). Each data point represents one biological replicate, consisting of the average of 2-3 technical replicates; horizontal bars indicate mean ± SEM. Comparison of total calcium transients between EtOH and ATc groups was analyzed using Fisher's exact test, $p$ < 0.0001 (see panel D').

**(D)** Total data from EtOH-treated (top row) parasites and ATc-treated (bottom row) parasites were categorized as in Supp. Fig. 3A-F within a 4×2 contingency table for comparison. Fisher's exact test was used for the comparison (right-most column). Each number represents the sum of 3 biological replicates, consisting of 2-3 technical replicates each.

**(D')** Data from (D) were summed based on the presence or absence of detected calcium transients (+ transient, –transient) and organized into 2×2 contingency tables. Fisher's exact test and odds ratio were used for the comparison.

**(E)** Comparison of odds ratios and effect sizes between the TATi parental line (D') and each of the TATi iKD lines (FER2, Nd9, NdP1, RASP2; Supp. Fig. 3C'-F', respectively.)

A

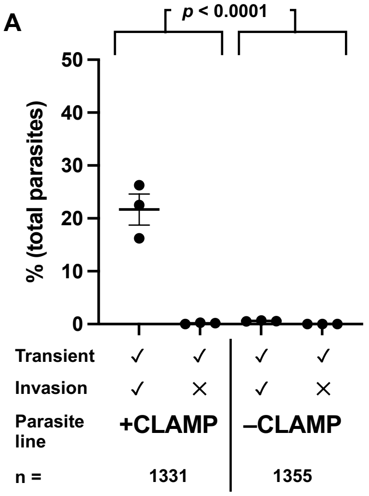


B **CLAMP**

| Treatment | +Transient +Invasion | +Transient –Invasion | –Transient +Invasion | –Transient –Invasion | Fisher's exact test |
|---|---|---|---|---|---|
| DMSO | 284 | 2 | 1 | 1044 | $p$ < 0.0001 |
| Rapamycin | 8 | 0 | 0 | 1347 | |

B'

| Treatment | +Transient | –Transient | Fisher's exact test |
|---|---|---|---|
| DMSO | 286 | 1045 | $p$ < 0.0001 |
| Rapamycin | 8 | 1347 | |

**Supp. Figure 6. Improved detection of calcium transients induced by parasites depleted of CLAMP protein.**

**(A)** Quantification of invasion events and calcium transients induced by control parasites (+CLAMP, n = 1331) compared to parasites depleted of CLAMP (-CLAMP, n = 1355) using a fluorescence detection method with increased sensitivity (Cal-520, probenecid treatment, 8 mM extracellular $CaCl_2$; see text for details). Each data point represents one biological replicate, consisting of the average of 2-3 technical replicates; horizontal bars indicate mean ± SEM. Comparison of total calcium transients between +CLAMP and -CLAMP groups was analyzed using Fisher's exact test, $p < 0.0001$ (Supp. Figure 6B').

**(B)** Total data from control-treated (top) parasites and CLAMP-depleted (bottom) parasites were categorized as in Supp. Fig. 3A-F within a 4×2 contingency table for comparison. Fisher's exact test was used for the comparison (right-most column). Each number represents the sum of 3 biological replicates, consisting of 3 technical replicates each.

**(B')** Data from (B) were summed based on the presence or absence of detected calcium transients (+ transient, –transient) and organized into 2×2 contingency tables. Fisher's exact test was used for the comparison (right-most column).

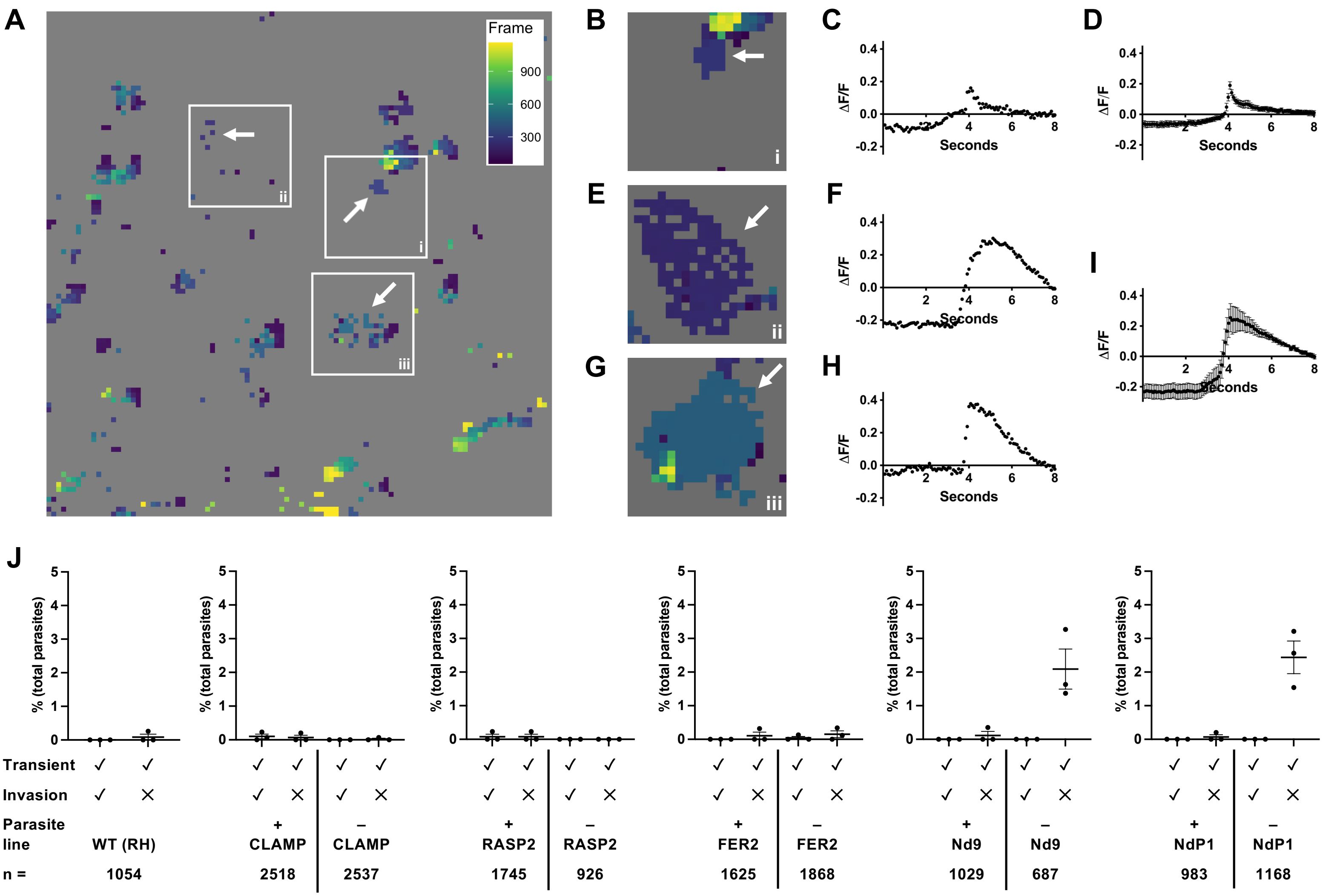


**Supp. Figure 7. Aberrant calcium transients detected in all parasite lines tested are typically associated with non-invading parasites.**

**(A)** Calcium transients generated by NdP1-depleted parasites. The PeakCaller output from the full 1020×1020 pixel FOV was plotted back to ROI location and color-coded by frame (inset) at which the transients reached their maximal intensity. Box i highlights a calcium transient associated with an invading parasite and boxes ii and iii highlight two aberrant calcium transients associated with parasites that subsequently failed to invade.

**(B, E, G)** Magnified views of the fluorescence signals detected in boxes i-iii from panel A, after reanalyzing the data with parameters optimized for capture of aberrant transient events.

**(C, F, G)** Quantification of Fluo-4 fluorescence levels (ΔF/F) in the host cell during the calcium transient events shown in panels B, E, G, respectively.

**(D, I)** Consensus plot of calcium transients generated by NdP1-depleted parasites that subsequently invaded the host cell (D, n = 15) and NdP1-depleted parasites that did not subsequently invade (I, n = 9). The fluorescence intensities in the 100 frames surrounding the peak of each calcium transient were averaged across all transients, the peaks of which were aligned to frame 51. The plot shows the mean ± SEM at each time point.

**(J)** Quantification of the frequency of aberrant spike generation by each of the parasite lines analyzed in this study. Each graph shows the number of aberrant spikes generated, as a percentage of the total number of parasites counted, and whether the aberrant spikes were associated with invading or non-invading parasites. Each data point represents one biological replicate, consisting of the average of 2-3 technical replicates. Horizontal bars indicate mean ± SEM.

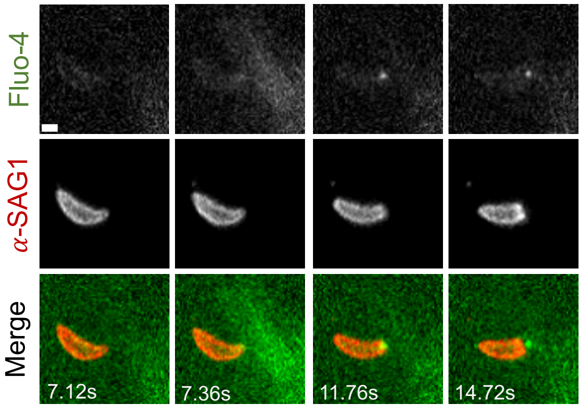


**Supp. Figure 8. Dot of Fluo-4 fluorescence at the apical end of a parasite invading a Fluo-4-loaded host cell.**

Individual frames from a time series showing changes in Fluo-4 fluorescence, including the appearance of a dot of fluorescence at the apical end of the parasite, and stripping of fluorescently-conjugated anti-SAG1 antibody from the surface of invading *T. gondii*. Top panels: Fluo-4 (pseudocolored green in merge); middle panels: anti-SAG1 (pseudocolored red in merge); bottom: merge. Scale bar = 2 µm. The calcium transient reaches maximal intensity at 7.36 seconds. Note that the dot of fluorescence moves past the moving junction and into the host cell along with the parasite (14.72 seconds), strongly suggesting that this fluorescent compartment is at or within the apical tip of the invading parasite. The full video from which these frames were extracted is presented as Supp. Video 3.
