## Supplemental video legends for "Perforation of the host cell plasma membrane during *Toxoplasma gondii* invasion requires rhoptry exocytosis"

### Supp. Video 1. A transient increase in intracellular calcium is observed within the host cell at the site of *T. gondii* invasion.

Video showing changes in Fluo-4 fluorescence (green) and stripping of fluorescently-conjugated anti-SAG1 antibody (red) from the surface of invading *T. gondii*. The calcium transient reaches maximal intensity at 6.32 seconds. Scale bar = 2 µm; time is shown as hr:min:sec.ms. Video shown at 1X speed (12.5 fps). Single frames from this video are shown in Fig. 2A.

### Supp. Video 2. NdP1-depleted parasite induces an aberrant calcium transient and fails to subsequently invade.

Video showing changes in Fluo-4 fluorescence (green) in the host cell during interaction with an NdP1-depleted parasite. Scale bar = 2 µm; time is shown as hr:min:sec.ms. Video shown at 4X speed (50 fps). Quantification of this calcium transient is shown in Supp. Fig. 7E-F.

### Supp. Video 3. Dot of Fluo-4 fluorescence at the apical end of a parasite invading a Fluo-4-loaded host cell.

Video showing changes in Fluo-4 fluorescence (green) within the host cell and the subsequent appearance of a dot of Fluo-4 fluorescence at the parasite apex during invasion. Scale bar = 2 µm; time is shown as hr:min:sec.ms. Video shown at 4X speed (50 fps). Single frames from this video are shown in Supp. Fig. 8.

### Supp. Video 4. Dot of Cal-520 fluorescence at the apical end of a parasite invading a Cal-520-loaded host cell – example 1.

Video showing changes in Cal-520 fluorescence (green) within the host cell and the subsequent appearance of a dot of Cal-520 fluorescence at the parasite apex during invasion. Scale bar = 2 µm; time is shown as hr:min:sec.ms. Video shown at 4X speed (50 fps).

### Supp. Video 5. Dot of Cal-520 fluorescence at the apical end of a parasite invading a Cal-520-loaded host cell – example 2.

Video showing changes in Cal-520 fluorescence (green) within the host cell and the subsequent appearance of a dot of Cal-520 fluorescence at the parasite apex during invasion. Scale bar = 2 µm; time is shown as hr:min:sec.ms. Video shown at 4X speed (50 fps).
